# Cross-sectional evaluation of humoral responses against SARS-CoV-2 Spike

**DOI:** 10.1101/2020.06.08.140244

**Authors:** Jérémie Prévost, Romain Gasser, Guillaume Beaudoin-Bussières, Jonathan Richard, Ralf Duerr, Annemarie Laumaea, Sai Priya Anand, Guillaume Goyette, Mehdi Benlarbi, Shilei Ding, Halima Medjahed, Antoine Lewin, Josée Perreault, Tony Tremblay, Gabrielle Gendron-Lepage, Nicolas Gauthier, Marc Carrier, Diane Marcoux, Alain Piché, Myriam Lavoie, Alexandre Benoit, Vilayvong Loungnarath, Gino Brochu, Elie Haddad, Hannah D. Stacey, Matthew S. Miller, Marc Desforges, Pierre J. Talbot, Graham T. Gould Maule, Marceline Côté, Christian Therrien, Bouchra Serhir, Renée Bazin, Michel Roger, Andrés Finzi

## Abstract

The SARS-CoV-2 virus is responsible for the current worldwide coronavirus disease 2019 (COVID-19) pandemic, infecting millions of people and causing hundreds of thousands of deaths. The Spike glycoprotein of SARS-CoV-2 mediates viral entry and is the main target for neutralizing antibodies. Understanding the antibody response directed against SARS-CoV-2 is crucial for the development of vaccine, therapeutic and public health interventions. Here we performed a cross-sectional study on 106 SARS-CoV-2-infected individuals to evaluate humoral responses against the SARS-CoV-2 Spike. The vast majority of infected individuals elicited anti-Spike antibodies within 2 weeks after the onset of symptoms. The levels of receptor-binding domain (RBD)-specific IgG persisted overtime, while the levels of anti-RBD IgM decreased after symptoms resolution. Some of the elicited antibodies cross-reacted with other human coronaviruses in a genus-restrictive manner. While most of individuals developed neutralizing antibodies within the first two weeks of infection, the level of neutralizing activity was significantly decreased over time. Our results highlight the importance of studying the persistence of neutralizing activity upon natural SARS-CoV-2 infection.

## MAIN

The first step in the replication cycle of coronaviruses is viral entry. This is mediated by their trimeric Spike (S) glycoproteins. Similar to SARS-CoV, the S glycoprotein of SARS-CoV-2 interacts with angiotensin-converting enzyme 2 (ACE2) as its host receptor (Hoffmann et al., 2020; Shang et al., 2020; Walls et al., 2019). During entry, the Spike binds the host cell through interaction between its receptor binding domain (RBD) and ACE2 and is cleaved by cell surface proteases or endosomal cathepsins (Hoffmann et al., 2020; Ou et al., 2020; Zang et al., 2020), triggering irreversible conformational changes in the S protein enabling membrane fusion and viral entry (Walls et al., 2020; Wrapp et al., 2020). The SARS-CoV-2 Spike is very immunogenic, with RBD representing the main target for neutralizing antibodies (Ju et al., 2020; Shi et al., 2020; Wu et al., 2020; Yuan et al., 2020). Humoral responses are important for preventing and controlling viral infections (Murin et al., 2019; Rouse and Sehrawat, 2010). However, little is known about the chronology and durability of the human antibody response against SARS-CoV-2.

Here we analyzed serological samples from 106 SARS-CoV-2-infected individuals at different times post-symptoms onset and 10 uninfected individuals for their reactivity to SARS-CoV-2 Spike glycoprotein, cross-reactivity with other human CoV (HCoV), as well as virus neutralization. Samples were collected from COVID-19 positive individuals starting on March 2020 or healthy individuals before the COVID-19 outbreak (COVID-19 negative). Cross-sectional serum samples (n=79) were collected from individuals presenting typical clinical symptoms of acute SARS-CoV-2 infection (Table 1). All patients were positive for SARS-CoV-2 by RT-PCR on nasopharyngeal specimens. The average age of the infected patients was 55 years old, including 33 males and 46 females. Samples were classified into 4 different time points after symptoms onset: 24 (11 males, 13 females) were obtained at 2-7 days (T1, median = 3 days), 20 (9 males, 11 females) between 8-14 days (T2, median = 11 days), 27 (10 males, 16 females) between 16-30 days (T3, median = 23 days) and 9 (3 males, 6 females) between 31-43 days (T4, median = 36 days). Samples were also obtained from 27 convalescent patients (20 males, 7 females, median = 41 days), who have been diagnosed with or tested positive for COVID-19 with complete resolution of symptoms for at least 14 days.

**Table 1.** Cross-sectional SARS-CoV-2 cohort clinical characteristics

| Group | n | Days after onset of symptoms<br>(median; day range) | Age<br>(median; age range) | Gender |  |
| --- | --- | --- | --- | --- | --- |
|  |  |  |  | Male (n) | Female (n) |
| T1 | 24 | 3<br>(2-7) | 50<br>(31-94) | 11 | 13 |
| T2 | 20 | 11<br>(8-14) | 64<br>(34-90) | 9 | 11 |
| T3 | 26 | 22<br>(16-30) | 40<br>(20-93) | 10 | 16 |
| T4 | 9 | 36<br>(31-43) | 39<br>(24-87) | 3 | 6 |
| Convalescent | 27 | 41<br>(23-52) | 37<br>(19-69) | 20 | 7 |

We first evaluated the presence of RBD-specific IgG and IgM antibodies by ELISA (Amanat et al., 2020; Stadlbauer et al., 2020). The level of RBD-specific IgM peaked at T2 and was followed by a stepwise decrease over time (T3, T4 and Convalescent) (Figure 1). Three quarter of the patients had detectable anti-RBD IgM two weeks after the onset of the symptoms. Similarly, 85% of patients in T2 developed anti-RBD IgG, reaching 100% in convalescent patients. In contrast to IgM, the levels of RBD-specific IgG peaked at T3 and remained relatively stable after complete resolution of symptoms (convalescent patients).

**Figure 1.**
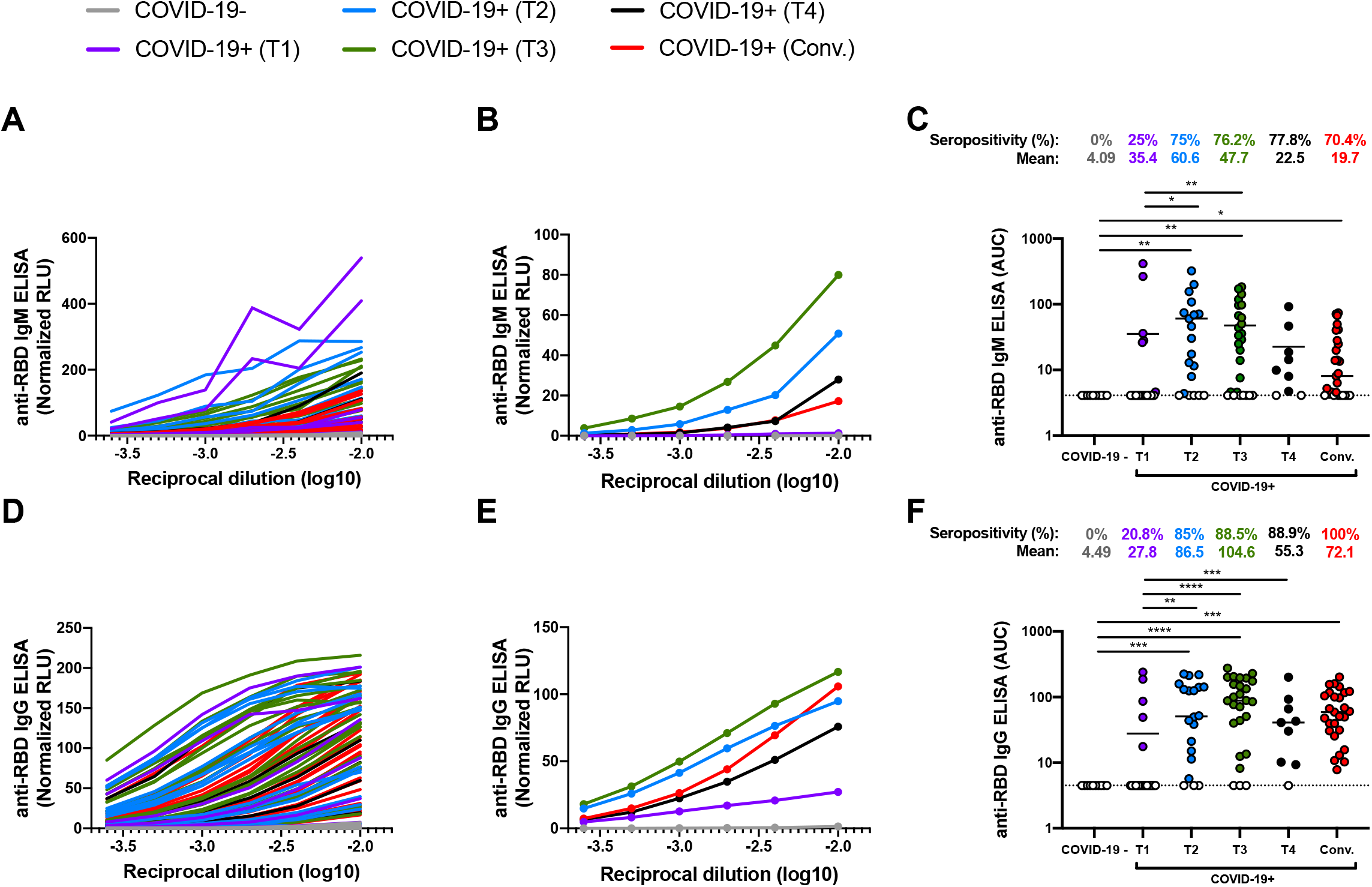
Detection of SARS-CoV-2 RBD-specific IgM and IgG over time. Indirect ELISA was performed using recombinant SARS-CoV-2 RBD and incubated with samples from COVID-19 negative or COVID-19 positive patients at different times after symptoms onset (T1, T2, T3, T4, Convalescent). Anti-RBD binding was detected using (A-C) anti-IgM-HRP or (D-F) anti-IgG-HRP. Relative light units (RLU) obtained with BSA (negative control) were subtracted and further normalized to the signal obtained with the anti-RBD CR3022 mAb present in each plate. Data in graphs (A, D) represent RLU done in quadruplicate. Curves depicted in (B, E) represent the mean RLU detected with all samples from the same group. Undetectable measures are represented as white symbols and limits of detection are plotted. (C, F) Areas under the curve (AUC) were calculated based on RLU datasets shown in (A, D) using GraphPad Prism software. Statistical significance was tested using Kruskal-Wallis tests with a Dunn’s post-test (* p < 0.05; ** p < 0.01; *** p < 0.001; **** p < 0.0001).

We next used flow cytometry to examine the ability of sera to recognize the full-length SARS-CoV-2 Spike expressed at the cell surface. Briefly, 293T cells expressing SARS-CoV-2 S glycoproteins were stained with samples, followed by incubation with secondary antibodies recognizing all antibody isotypes (including IgG, IgM and IgA). As presented in Figure 2, 54.2% of the sera from T1 already contained SARS-CoV-2 full Spike-reactive antibodies. Interestingly, the majority of patients from T2, T3, T4 and convalescent groups were found to be seropositive in agreement with previous report (Grzelak et al., 2020). The higher seropositivity detected by flow cytometry is most likely due to the detection of antibodies of multiple specificity and of different isotypes simultaneously. Antibody levels targeting the SARS-CoV-2 Spike significantly increased from T1 to T2/T3 and remained relatively stable thereafter. As expected, the levels of antibodies recognizing the full Spike correlated with the presence of both RBD-specific IgG and IgM (Figure S1). We also evaluated potential cross-reactivity against the closely related SARS-CoV Spike. None of the COVID-19 negative samples recognized the SARS-CoV Spike. While the reactivity of COVID-19+ samples to SARS-CoV S was lower than for SARS-CoV-2 S, it followed a similar progression and significantly correlated with their reactivity to SARS-CoV-2 full Spike or RBD protein (Figure 2 and S1). This indicates that SARS-CoV-2-elicited antibodies cross-react with human *Sarbecoviruses*. This was also observed with another *Betacoronavirus* (OC43) but not with *Alphacoronavirus* (NL63, 229E) S glycoproteins, suggesting a genus-restrictive cross-reactivity (Figure 2C and S1). Of note, anti-OC43 RBD antibodies did not fluctuate upon SARS-CoV-2 infection (Figure S2). Therefore, this differential cross-reactivity could be explained by the high degree of conservation in the S protein fusion machinery, particularly in the S2 subunit among *Betacoronaviruses* (Jaimes et al., 2020; Madu et al., 2009; Zhou et al., 2020).

**Figure 2.**
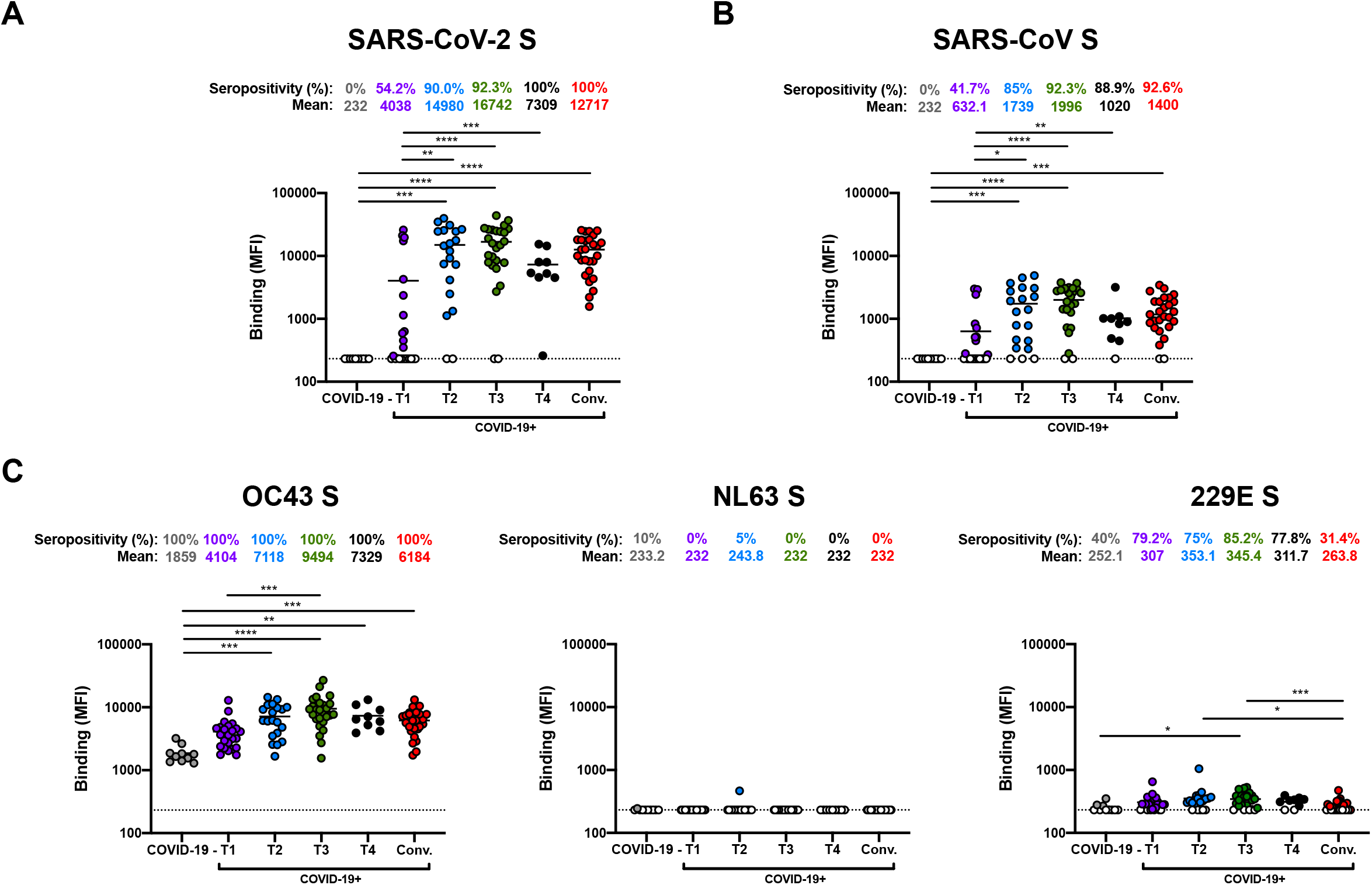
SARS-CoV-2 infection elicits cross-reactive antibodies against other human *Betacoronaviruses*. Cell-surface staining of 293T cells expressing full-length Spike (S) from different HCoV (A) SARS-CoV-2, (B) SARS-CoV, (C) OC43, NL63 and 229E with samples from COVID-19 negative or COVID-19 positive patients at different stage of infection (T1, T2, T3, T4, Convalescent). The graphs shown represent the median fluorescence intensities (MFI). Undetectable measures are represented as white symbols and limits of detection are plotted. Error bars indicate means ± SEM. Statistical significance was tested using Kruskal-Wallis tests with a Dunn’s post-test (* p < 0.05; ** p < 0.01; *** p < 0.001; **** p < 0.0001).

We next measured the capacity of patient samples to neutralize pseudoparticles bearing SARS-CoV-2 S, SARS-CoV S or VSV-G glycoproteins using 293T cells stably expressing ACE2 as target cells (Figure 3 and S3). Neutralizing activity, as measured by the neutralization half-maximum inhibitory dilution (ID_50_) or the neutralization 80% inhibitory dilution (ID_80_), was detected in most patients within 2 weeks after the onset of symptoms (T2, T3, T4 and Convalescent patients) (Figure 3). SARS-CoV-2 neutralization was specific since no neutralization was observed against pseudoparticles expressing VSV-G. The capacity to neutralize SARS-CoV-2 S-pseudotyped particles significantly correlated with the presence of RBD-specific IgG/IgM and anti-S antibodies (Figure S4). While the percentage of patients eliciting neutralizing antibodies against SARS-CoV-2 Spike remained relatively stable 2 weeks after disease symptom onset (T2, T3, T4 and Convalescent patients), neutralizing antibody titers significantly decreased after 1 month of infection (T4) or after the complete resolution of symptoms as observed in the convalescent patients (Figure 3G and 3H). Similarly to RBD-specific IgM, levels of RBD-specific IgA were also found to peak at T2 and decrease over time. However, RBD-specific IgM levels displayed a stronger correlation with neutralization acitivity compared to RBD-specific IgG and IgA, suggesting a more prominent role for IgM, but the decrease in IgA could also contribute to the loss of neutralization activity as recently suggested (Sterlin et al., 2020). Cross-reactive neutralizing antibodies against SARS-CoV S protein (Figure 2B) were also detected in some SARS-CoV-2-infected individuals, but with significantly lower potency and waned over time. We note that around 40% of convalescent patients did not exhibit any neutralizing activity. This suggests that the production of neutralizing antibodies is not a prerequisite to the resolution of the infection and that other arms of the immune system could be sufficient to control the infection in an important proportion of the population.

**Figure 3.**
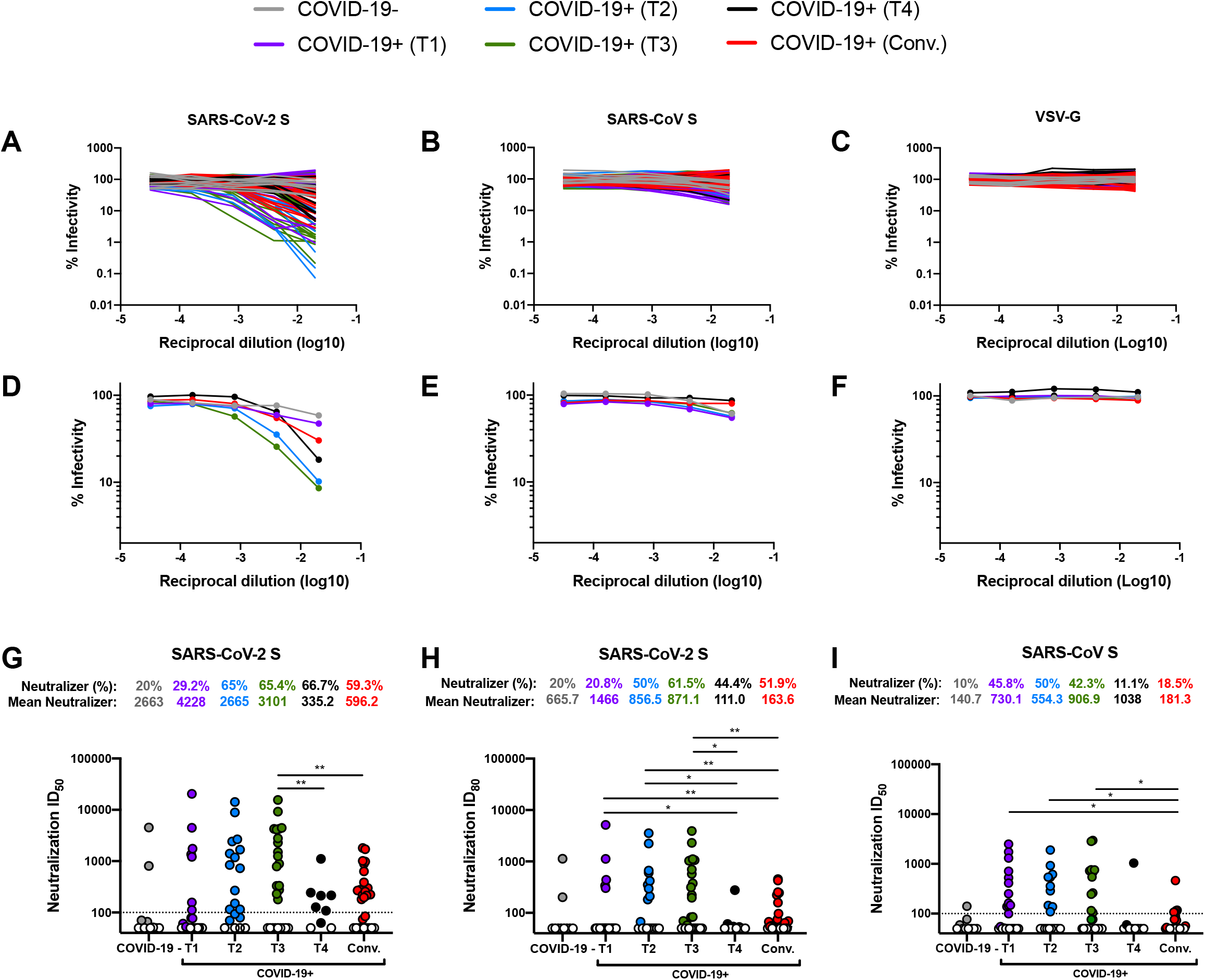
Anti-Spike neutralizing antibody titers decrease over time. Pseudoviral particles coding for the luciferase reporter gene and bearing the following glycoproteins: (A, D, G, H) SARS-CoV-2 S, (B, E, I) SARS-CoV S or (C, F) VSV-G were used to infect 293T-ACE2 cells. Pseudoviruses were incubated with serial dilutions of samples from COVID-19 negative or COVID-19 positive patients (T1, T2, T3, T4, Convalescent) at 37°C for 1h prior to infection of 293T-ACE2 cells. Infectivity at each dilution was assessed in duplicate and is shown as the percentage of infection without sera for each glycoprotein. (G, I) Neutralization half maximal inhibitory serum dilution (ID_50_) and (H) ID_80_ values were determined using a normalized non-linear regression using Graphpad Prism software. Undetectable measures are represented as white symbols. Neutralizer represent patients with (G, I) an ID_50_ over 100 or (H) an ID_80_. Statistical significance was tested using Mann-Whitney U tests (* p < 0.05; ** p < 0.01).

To determine whether underlying correlation patterns among antibody responses detected in SARS-CoV-2 infected individuals were associated with demographic and clinical parameters, we performed a comprehensive correlation analysis, focusing on data from the acute stages of SARS-CoV-2 infection (T1, T2, T3 and T4) (Figure 4 & S5). This analysis revealed a prominent cluster of positive correlations between SARS-CoV-2, SARS-CoV, and OC43 Spike antibody binding, SARS-CoV-2 neutralization, and days post-symptoms onset (Figure S5). The cluster became evident in a linear correlation analysis involving all study parameters (Figure S5A). Of interest, clinical parameters formed another cluster of positive correlations between respiratory symptoms, hospitalization, oxygen supplementation and intensive care unit (ICU) admission (Figure S5A). The presence of respiratory symptoms and hospitalization also correlated with age of the infected patients. Studying the network of immunologic and clinical correlation pairs longitudinally (from T1 to T4), we observed an increased diversification of associations between the parameters (Figure 4B-E), Associations between anti-Spike Abs and clinical parameters enhanced overtime and was more prominent 3 weeks after the onset of the symptoms (T3 & T4). Admission to the ICU was significantly associated with levels of RBD-specific IgM and IgG and total SARS-CoV-2 Spike Abs (Figure 4A & S5A). The presence of respiratory symptoms was linked to higher levels of RBD-specific IgM and of neutralization activity against SARS-CoV-2 S (Figure 4A). Indeed, neutralizers (patients with detectable neutralization ID_50_ against SARS-CoV-2) were found to have stronger antibody responses and were more inclined to present respiratory symptoms (Figure S6).

**Figure 4.**
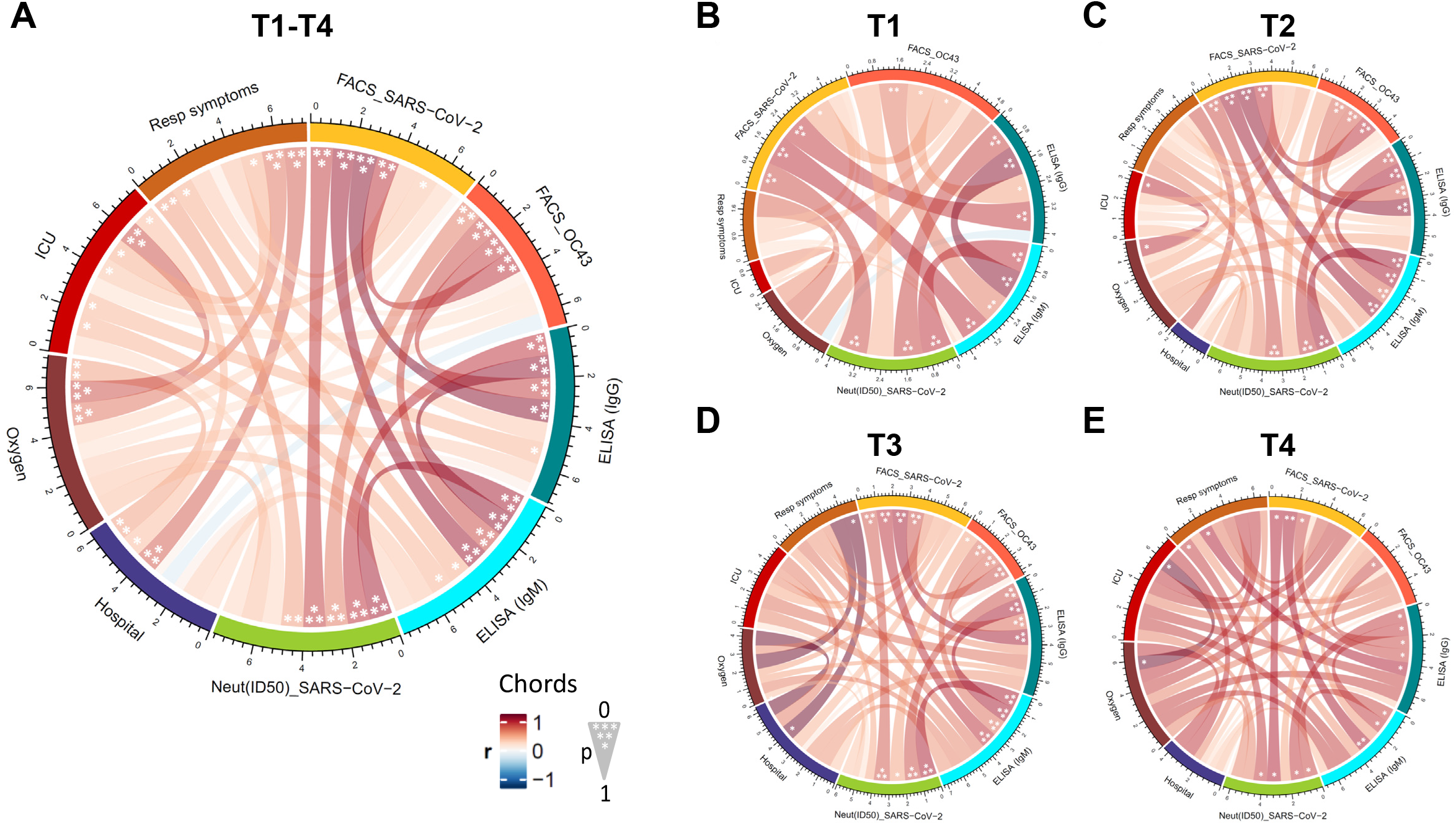
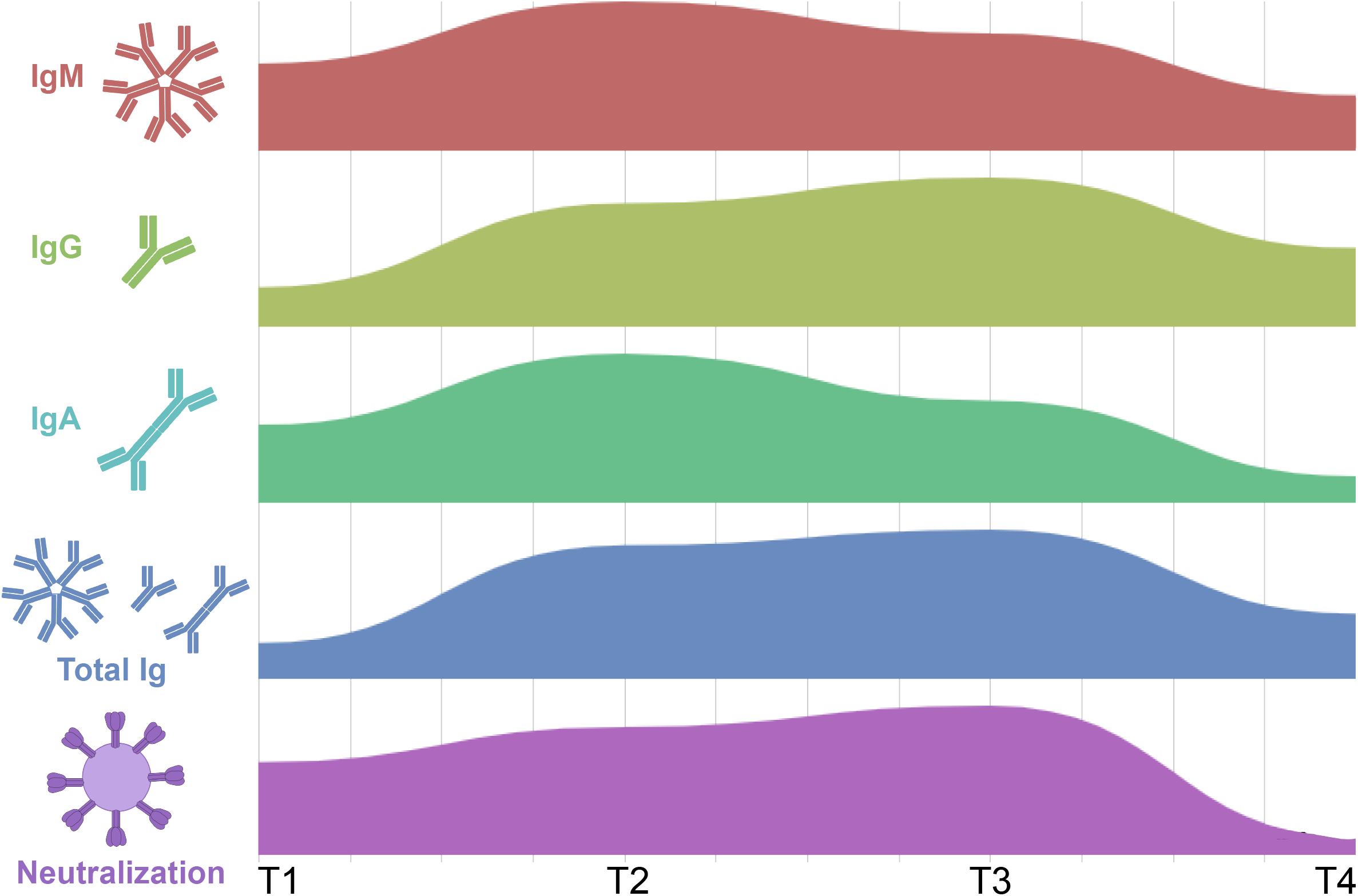
Association between clinical and serological parameters in SARS-CoV-2-infected patients. Chord diagram illustrating the network of linear correlations among nine major serological and clinical factors for (A) all acutely infected individuals (T1, T2, T3 and T4) or (B-E) at different time points. Chords are color-coded according to the magnitude of the correlation coefficient (r); chord width inversely corresponds to the P-value. Asterisks indicate all statistically significant correlations within chords (*P < 0.05, **P < 0.01, ***P < 0.005). (A-E) Correlation analysis was done using nonparametric Spearman rank tests. P-values were adjusted for multiple comparisons using Holm-Sidak (α = 0.05). Statistical comparisons of two parameters were done using Mann-Whitney U tests.

This study helps to better understand the kinetics and persistence of humoral responses directed against SARS-CoV-2 (Figure 1, 2 & 3). Our results reveal that the vast majority of infected individuals are able to elicit antibodies directed against SARS-CoV-2 Spike within 2 weeks after symptom onset and persist after the resolution of the infection. Accordingly, all tested convalescent patients were found to be seropositive. As expected, RBD-specific IgM levels decreased over the duration of the study while IgG remained relatively stable. Our results highlight how SARS-CoV-2 Spike, like other coronaviruses, appears to be relatively easily recognized by Abs present in sera from infected individuals. This was suggested to be linked to the higher processing of glycans compared to other type I fusion protein, such as HIV-1 Env, Influenza A HA or filoviruses GP (Watanabe et al., 2020a; Watanabe et al., 2020b). The ease of naturally-elicited Abs to recognize the Spike might be associated with the low rate of somatic hypermutation observed in neutralizing Abs (Ju et al., 2020). This low somatic hypermutation rate could in turn explain why the majority of the SARS-CoV-2 infected individuals are able to generate neutralizing antibodies within only two weeks after infection (Figure 3). In contrast, the development of potent neutralizing antibodies against HIV-1 Env usually requires 2-3 years of infection and require a high degree of somatic hypermutation (Sok and Burton, 2018). Nevertheless, in the case of SARS-CoV-2 infection, the neutralization capacity decreases significantly 6 weeks after the onset of symptoms, following a similar trend as anti-RBD IgM (Figure 1 & 3). Interestingly, anti-RBD IgM presented a stronger correlation with neutralization than IgG and IgA (Figure S4A,C), suggesting that at least part of the neutralizing activity is mediated by IgM. The neutralization activity appears to further decrease after the resolution of symptoms as recently reported in a series of longitudinal studies on convalescent patients (Beaudoin-Bussières et al., 2020; Ibarrondo et al., 2020; Long et al., 2020; Perreault et al., 2020; Yin et al., 2020; Zhang et al., 2020). However, it remains unclear whether this reduced level of neutralizing activity would remain sufficient to protect from re-infection.

## AUTHOR CONTRIBUTIONS

J.Prévost, J.R., B.S., R.B., M.R. and A.F. conceived the study. J.Prévost, J.R., A.F. designed experimental approaches; J.Prévost, G.B.B., R.G., A.Laumaea, J.R., S.P.A., G.G., M.B., S.D., T.T., J.Perreault, A.Lewin., R.D. R.B., M.R., and A.F. performed, analyzed and interpreted the experiments; J.Prévost, G.B.B., J.R., H.M., G.G.-L., H.D.S., M.S.M., M.D., P.T., G.T.G.M., M.Côté and A.F. contributed novel reagents; N.G., M.Carrier, D.M., A.P., M.L., A.B., V.L., G.B., E.H., C.T., R.B. and M.R. collected clinical samples; J.Prévost, J.R. and A.F. wrote the paper. Every author has read edited and approved the final manuscript.

## ACKNOWLEDGMENTS

The authors thank the CRCHUM BSL3 and Flow Cytometry Platforms for technical assistance. We thank Dr Florian Krammer (Icahn School of Medicine at Mount Sinai, NY) for the plasmid expressing the SARS-CoV-2 RBD domain, Dr Stefan Pöhlmann (Georg-August University, Germany) for the plasmids coding for SARS-CoV S, SARS-CoV-2 S and HCoV 229E and NL63 S glycoproteins and Dr M. Gordon Joyce (U.S. MHRP) for the monoclonal antibody CR3022. We also thank Danka K Shank and Melina Bélanger Collard from the Laboratoire de Santé Publique du Québec for their help in preparing the specimens. This work was supported by le Ministère de l’Économie et de l’Innovation du Québec, Programme de soutien aux organismes de recherche et d’innovation to A.F and by the Fondation du CHUM. This work was also supported by a CIHR foundation grant #352417 to A.F. Development of SARS-CoV-2 reagents was partially supported by the NIAID Centers of Excellence for Influenza Research and Surveillance (CEIRS) contract HHSN272201400008C. A.F. is the recipient of a Canada Research Chair on Retroviral Entry # RCHS0235 950-232424. R.D. was supported by NIH grant R01 AI122953-05. M.C. is the recipient of a Tier II Canada Research Chair in Molecular Virology and Antiviral Therapeutics and an Ontario’s Early Researcher Award. J.P., G.B.B. and S.P.A are supported by CIHR fellowships. R.G. is supported by a MITACS Accélération postdoctoral fellowship. M.S.M was supported in part by a CIHR New Investigator Award, and Ontario Early Researcher Award, CIHR COVID Rapid Response Funding, and the W. Garfield Weston Foundation Weston Family Microbiome Initiative. The funders had no role in study design, data collection and analysis, decision to publish, or preparation of the manuscript.

## COMPETING INTERESTS

The authors declare no competing interests.

## STAR METHODS

### Ethics statement

All work was conducted in accordance with the Declaration of Helsinki in terms of informed consent and approval by an appropriate institutional board. In addition, this study was conducted in accordance with the rules and regulations concerning ethical reviews in Quebec, particularly those specified in the Civil Code (http://legisquebec.gouv.qc.ca/fr/ShowDoc/cs/CCQ-1991) and in subsequent IRB practice. Informed Consent was obtained for all participating subjects and the study was approved by Quebec Public health authorities. Convalescent plasmas were obtained from donors who consented to participate in this research project (REB # 2020-004). The donors were recruited by Héma-Québec and met all donor eligibility criteria for routine apheresis plasma donation, plus two additional criteria: previous confirmed COVID-19 infection and complete resolution of symptoms for at least 14 days. Plasma samples from COVID-children were obtained from donors enrolled in a research protocol from CHU Ste-Justine (REB #3195).

### Plasmids

The plasmids expressing the human coronavirus Spikes of SARS-CoV-2, SARS-CoV, NL63 and 229E were previously reported (Hoffmann et al., 2020; Hofmann et al., 2005). The OC43 Spike with an N-terminal 3xFlag tag and C-terminal 17 residue deletion was cloned into pCAGGS following amplification of the spike gene from pB-Cyst-3FlagOC43SC17 (kind gift of James M. Rini, University of Toronto, ON, Canada). The plasmid encoding for SARS-CoV-2 S RBD (residues 319-541) fused with a hexahistidine tag was reported elsewhere (Amanat et al., 2020). The sequence for the HCoV OC43 RBD was obtained from the UniProt Protein Database (P36334 SPIKE_CVHOC). An N-terminal 13aa signal sequence and a C-terminal His-tag were added for downstream protein purification. Mammalian cell codon optimization was performed using the GenScript GenSmart Codon Optimization Tool. The RBD gene was synthesized by GenScript and cloned into the pcDNA3.1 plasmid between EcoRI and XhoI sites. The vesicular stomatitis virus G (VSV-G)-encoding plasmid (pSVCMV-IN-VSV-G) was previously described (Lodge et al., 1997). The lentiviral packaging plasmids pLP1 and pLP2, coding for HIV-1 gag/pol and rev respectively, were purchased from Invitrogen. The transfer plasmid (pLenti-C-mGFP-P2A-Puro-ACE2) encoding for human angiotensin converting enzyme 2 (ACE2) fused with a mGFP C-terminal tag and a puromycin selection marker was purchased from OriGene.

### Cell lines

293T human embryonic kidney cells (obtained from ATCC) were maintained at 37°C under 5% CO_2_ in Dulbecco’s modified Eagle’s medium (DMEM) (Wisent) containing 5% fetal bovine serum (VWR) and 100 μg/ml of penicillin-streptomycin (Wisent). For the generation of 293T cells stably expressing human ACE2, transgenic lentivirus were produced in 293T using a third-generation lentiviral vector system. Briefly, 293T cells were co-transfected with two packaging plasmids (pLP1 and pLP2), an envelope plasmid (pSVCMV-IN-VSV-G) and a lentiviral transfer plasmid coding for human ACE2 (pLenti-C-mGFP-P2A-Puro-ACE2) (OriGene). Forty-eight hours post-transfection, supernatant containing lentiviral particles was used to infect more 293T cells in presence of 5μg/mL polybrene. Stably transduced cells were enriched upon puromycin selection. 293T-ACE2 cells were then cultured in a medium supplemented with 2 μg/ml of puromycin (Sigma).

### Protein expression and purification

FreeStyle 293F cells (Invitrogen) were grown in FreeStyle 293F medium (Invitrogen) to a density of 1 x 106 cells/mL at 37°C with 8 % CO2 with regular agitation (150 rpm). Cells were transfected with a plasmid coding for SARS-CoV-2 S RBD or OC43 S RBD using ExpiFectamine 293 transfection reagent, as directed by the manufacturer (Invitrogen). One week later, cells were pelleted and discarded. Supernatants were filtered using a 0.22 μm filter (Thermo Fisher Scientific). The recombinant RBD proteins were purified by nickel affinity columns, as directed by the manufacturer (Invitrogen). The RBD preparations were dialyzed against phosphate-buffered saline (PBS) and stored in aliquots at −80°C until further use. To assess purity, recombinant proteins were loaded on SDS-PAGE gels and stained with Coomassie Blue. For cell-surface staining, RBD proteins were fluorescently labelled with Alexa Fluor 594 (Invitrogen) according to the manufacturer’s protocol.

### Sera and antibodies

Sera from SARS-CoV-2-infected and uninfected donors were collected, heat-inactivated for 1 hour at 56 °C and stored at −80°C until ready to use in subsequent experiments. The monoclonal antibodies CR3022 and 4.3E4 were used as positive controls in ELISA assays and were previously described (Desforges et al., 2013; ter Meulen et al., 2006; Tian et al., 2020; Yuan et al., 2020). Horseradish peroxidase (HRP)-conjugated antibody specific for the Fc region of human IgG (Invitrogen), for the Fc region of human IgM (Jackson ImmunoReasearch) or for the Fc region of human IgA (Jackson ImmunoResearch) were used as secondary antibodies to detect sera binding in ELISA experiments. Alexa Fluor-647-conjugated goat anti-human IgG (H+L) Abs (Invitrogen) were used as secondary antibodies to detect sera binding in flow cytometry experiment. Polyclonal goat anti-ACE2 (R&D systems) and Alexa-Fluor-conjugated donkey anti-goat IgG Abs (Invitrogen) were used to detect cell-surface expression of human ACE2.

### ELISA assay

Recombinant SARS-CoV-2 S RBD proteins (or OC43 S RBD proteins) (2.5 □ μg/ml), or bovine serum albumin (BSA) (2.5 □ μg/ml) as a negative control, were prepared in PBS and were adsorbed to plates (MaxiSorp; Nunc) overnight at 4°C. Coated wells were subsequently blocked with blocking buffer (Tris-buffered saline [TBS] containing 0.1% Tween20 and 2% BSA) for 1h at room temperature. Wells were then washed four times with washing buffer (Tris-buffered saline [TBS] containing 0.1% Tween20). CR3022 mAb (50ng/ml) or sera from SARS-CoV-2-infected or uninfected donors (1/100; 1/250; 1/500; 1/1000; 1/2000; 1/4000) were diluted in blocking buffer and incubated with the RBD-coated wells for 1h at room temperature. Plates were washed four times with washing buffer followed by incubation with secondary Abs (diluted in blocking buffer) for 1h at room temperature, followed by four washes. HRP enzyme activity was determined after the addition of a 1:1 mix of Western Lightning oxidizing and luminol reagents (Perkin Elmer Life Sciences). Light emission was measured with a LB941 TriStar luminometer (Berthold Technologies). Signal obtained with BSA was subtracted for each serum and were then normalized to the signal obtained with CR3022 mAb present in each plate. Alternatively, the signal obtained with each serum on OC43 RBD was normalized with the signal obtained with 4.3E4 mAb present in each plate. The seropositivity threshold was established using the following formula: mean RLU of all COVID-19 negative sera normalized to CR3022 (or 4.3E4) + (3 standard deviations of the mean of all COVID-19 negative sera).

### Flow cytometry analysis of cell-surface staining

Using the standard calcium phosphate method, 10μg of Spike expressor and 2μg of a green fluorescent protein (GFP) expressor (pIRES-GFP) was transfected into 2 × 10^6^ 293T cells. At 48h post transfection, 293T cells were stained with sera from SARS-CoV-2-infected or uninfected individuals (1:250 dilution). The percentage of transfected cells (GFP+ cells) was determined by gating the living cell population based on the basis of viability dye staining (Aqua Vivid, Invitrogen). Samples were acquired on a LSRII cytometer (BD Biosciences, Mississauga, ON, Canada) and data analysis was performed using FlowJo vX.0.7 (Tree Star, Ashland, OR, USA). The seropositivity threshold was established using the following formula: (mean of all COVID-19 negative sera + (3 standard deviation of the mean of all COVID-19 negative sera) + inter-assay coefficient of variability).

### Virus neutralization assay

Target cells were infected with single-round luciferase-expressing lentiviral particles. Briefly, 293T cells were transfected by the calcium phosphate method with the lentiviral vector pNL4.3 R-E-Luc (NIH AIDS Reagent Program) and a plasmid encoding for SARS-CoV-2 Spike, SARS-CoV Spike or VSV-G at a ratio of 5:4. Two days post-transfection, cell supernatants were harvested and stored at −80°C until use. 293T-ACE2 target cells were seeded at a density of 1×10^4^ cells/well in 96-well luminometer-compatible tissue culture plates (Perkin Elmer) 24h before infection. Recombinant viruses in a final volume of 100μl were incubated with the indicated sera dilutions (1/50; 1/250; 1/1250; 1/6250; 1/31250) for 1h at 37°C and were then added to the target cells followed by incubation for 48h at 37°C; cells were lysed by the addition of 30μl of passive lysis buffer (Promega) followed by one freeze-thaw cycle. An LB941 TriStar luminometer (Berthold Technologies) was used to measure the luciferase activity of each well after the addition of 100 μl of luciferin buffer (15mM MgSO_4_, 15mM KPO_4_ [pH 7.8], 1mM ATP, and 1mM dithiothreitol) and 50μl of 1mM d-luciferin potassium salt (Prolume). The neutralization half-maximal inhibitory dilution (ID_50_) or the neutralization 80% inhibitory dilution (ID_80_) represents the sera dilution to inhibit 50% or 80% of the infection of 293T-ACE2 cells by recombinant viruses bearing the indicated surface glycoproteins.

### Software Scripts and Visualization

Correlograms were generated using the corrplot package in program R and R Studio (R Core Team, 2013; R Studio Team, 2015). Dendrograms were calculated using the dendPlot function and hclust method, or as implemented in the heatmap package in R. Chord diagrams were generated in R and R Studio based on the circlize and ComplexHeatmap package, as recently described. For time series, area graphs were generated using RawGraphs with DensityDesign interpolation and the implemented normalization using vertically un-centered values (Mauri et al., 2017). Forrest plots and calculations of fold change, significance (Mann-Whitney) and adjusted P values (Holm-Sidak) were done using Excel and Prism v8.2.0. The confidence interval of a quotient of two means was calculated based on the Fieller method using GraphPad QuickCalcs.

### Statistical analyses

Statistics were analyzed using GraphPad Prism version 8.0.2 (GraphPad, San Diego, CA, (USA). Every data set was tested for statistical normality and this information was used to apply the appropriate (parametric or nonparametric) statistical test. P values <0.05 were considered significant; significance values are indicated as * P<0.05, ** P<0.01, *** P<0.001, **** P<0.0001. Corrections for multiple comparisons were performed with the Holm-Sidak method.

## Supplemental Information

Supplemental information includes 1 table and 6 figures, and can be found online.

**Supplemental Table 1.**
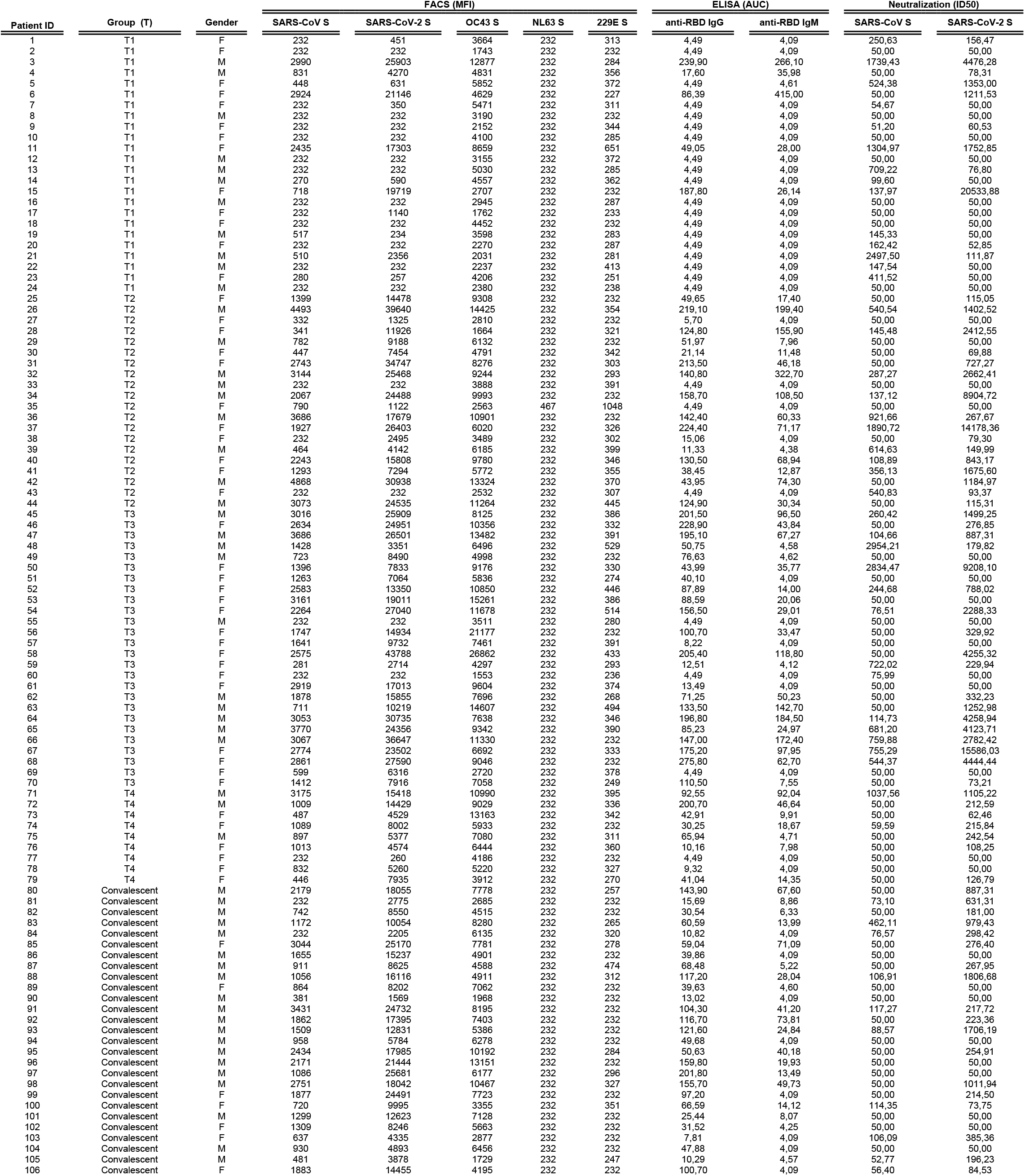
Serological analysis of samples from SARS-CoV-2 infected individuals

**Supplemental Figure 1.**
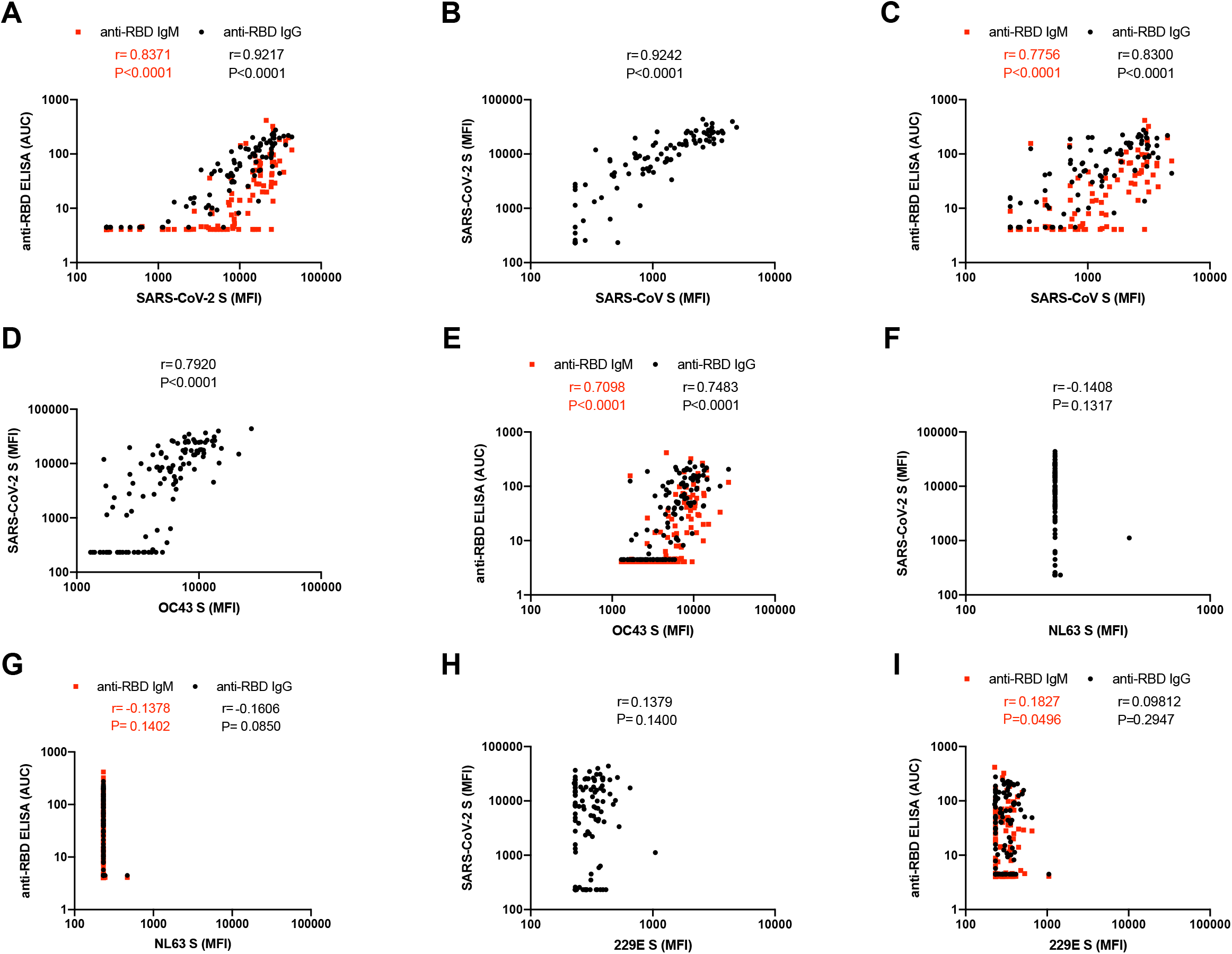
Detection of antibodies against cell-surface expressed SARS-CoV-2 full Spike correlates with RBD-specific IgG and IgM. (A,C,E,G,I) Levels of recognition of the different human coronavirus Spikes (SARS-CoV-2 S, SARS-CoV, OC43 S, NL63 S, 229E S) evaluated by flow cytometry (Figure 2) were plotted against the levels of anti-RBD IgG and IgM evaluated by indirect ELISA (Figure 1). (B,D,F,H) Levels of recognition of different HCoV Spikes (SARS-CoV, OC43 S, NL63 S, 229E S) evaluated by flow cytometry were plotted against the levels of recognition of SARS-CoV-2 S (also evaluated by flow cytometry). Statistical analysis was performed using Spearman rank correlation tests.

**Supplemental Figure 2.**
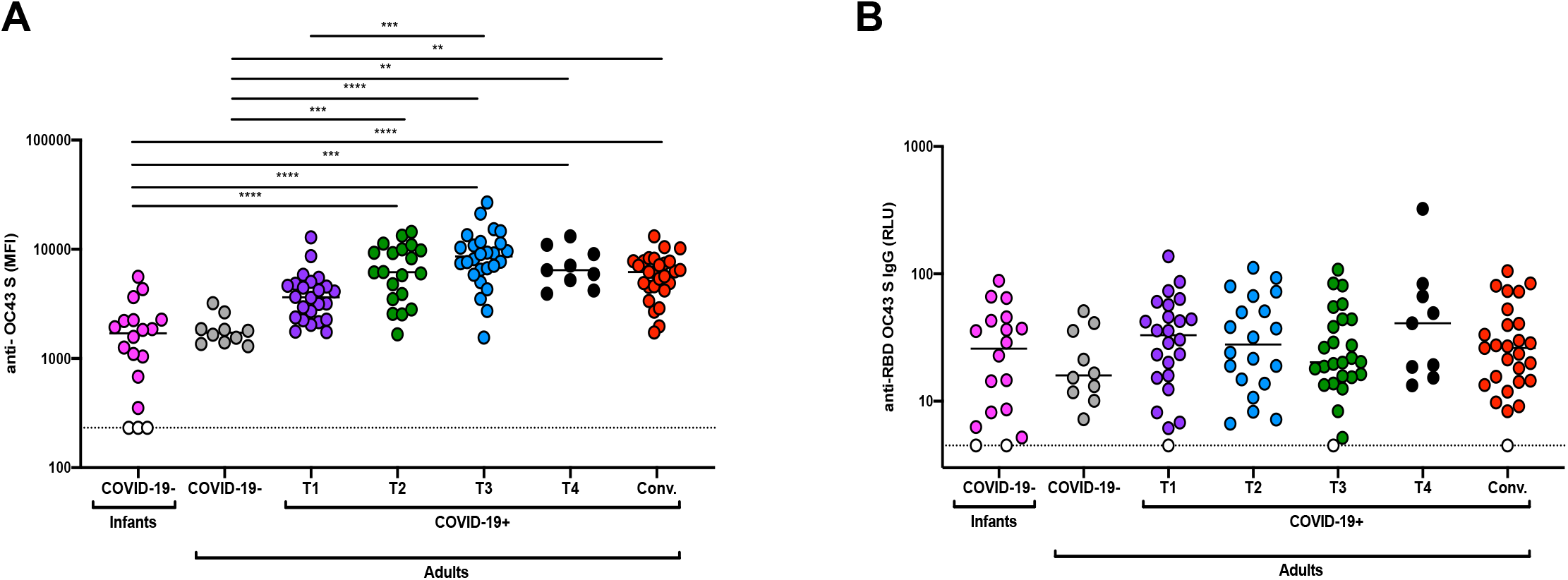
Time course of antibodies against OC43 Spike upon SARS-CoV-2 infection. (A) Cell-surface staining of 293T cells expressing full-length OC43 Spike (S) and (B) indirect ELISA using recombinant OC43 RBD. S-expressing cells or RBD-coated wells were incubated with samples from COVID-19 negative infants and adults or COVID-19 positive patients at different times after symptoms onset (T1, T2, T3, T4, Convalescent). (A) The graphs shown represent the median fluorescence intensities (MFI). Undetectable measures are represented as white symbols and limits of detection are plotted. (B) Anti-RBD binding was detected using anti-IgG-HRP. Relative light units (RLU) obtained with BSA (negative control) were subtracted and further normalized to the signal obtained with the anti-OC43 RBD 4.3E4 mAb present in each plate. Data in graphs represent RLU done in quadruplicate, with error bars indicating means ± SEM. Undetectable measures are represented as white symbols and limits of detection are plotted. Statistical significance was tested using Kruskal-Wallis tests with a Dunn’s post-test (** P < 0.01; *** P < 0.001; **** P < 0.0001).

**Supplemental Figure 3.**
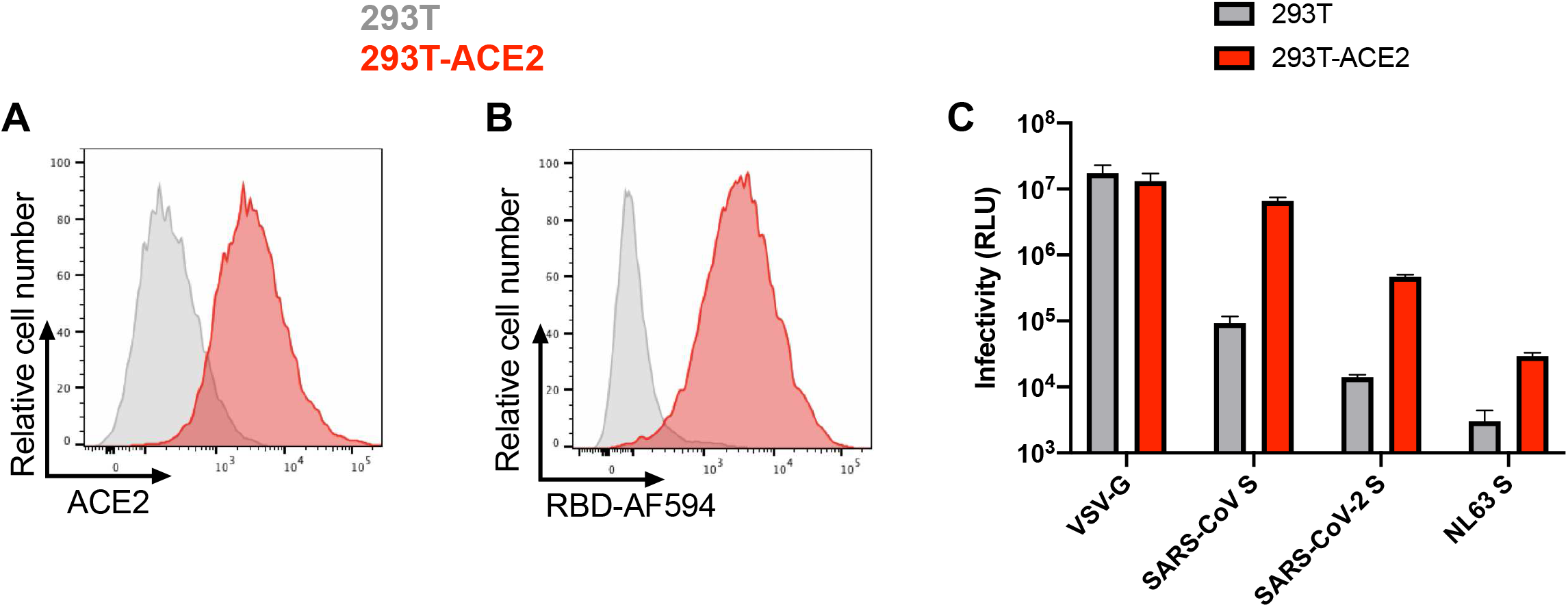
Characterization of 293T-ACE2 cell line. Cell-surface staining of 293T cells and 293T stably expressing human ACE2 (293T-ACE2) with (A) polyclonal goat anti-ACE2 or (B) RBD conjugated with Alexa Fluor 594 (RBD-AF594). Shown in (A,B) are histograms depicting representative anti-ACE2 and RBD-AF594 staining. (C) Recombinant pseudovirus expressing luciferase and bearing SARS-CoV-2 or VSV-G glycoproteins were used to infect 293T or 293T-ACE2 and infectivity was quantified by luciferase activity in cell lysate by relative light units (RLU).

**Supplemental Figure 4.**
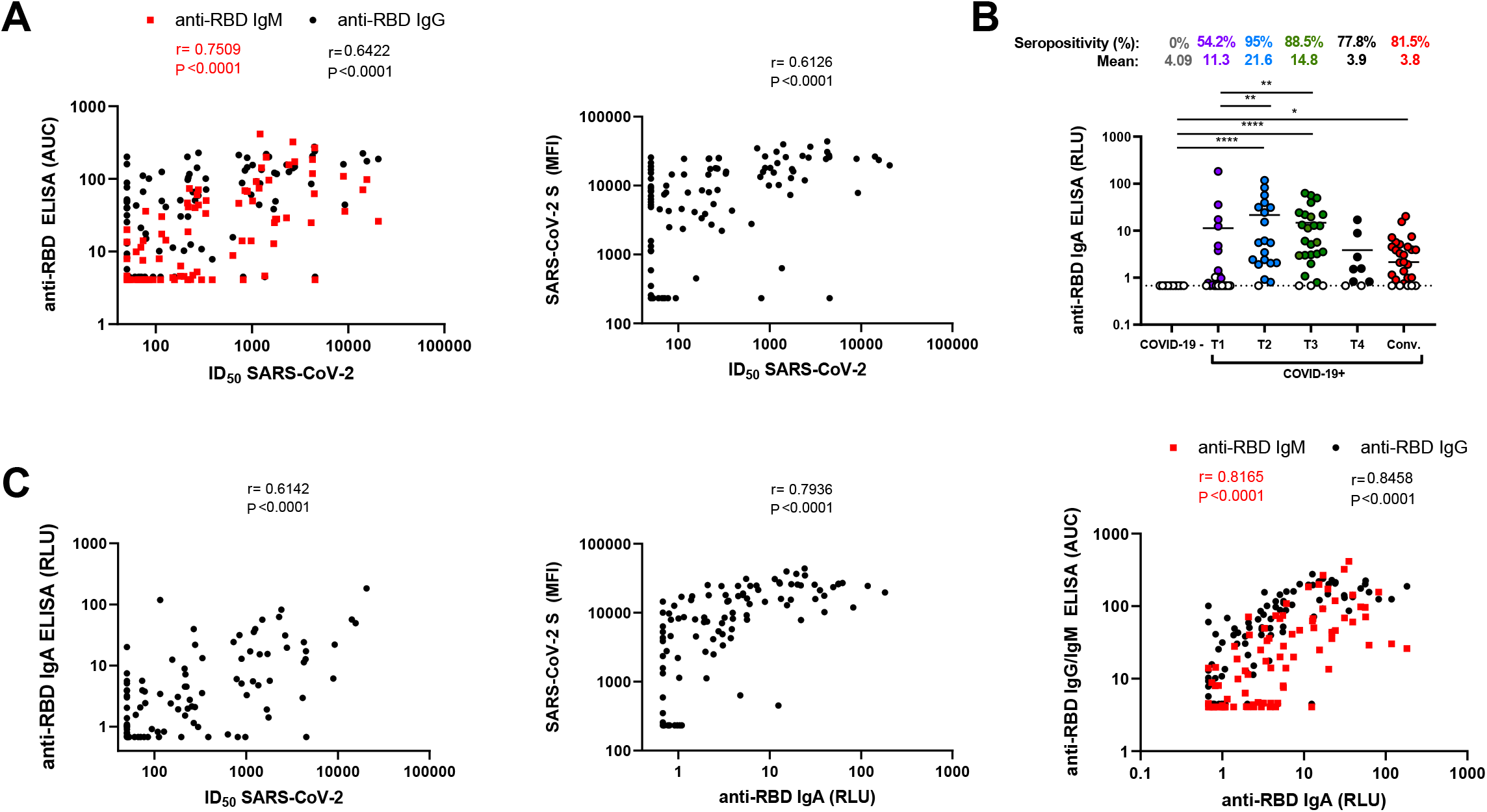
Anti-RBD antibodies positively correlate with neutralization. (A) The neutralization ID_50_ with SARS-CoV-2 S was correlated with the levels of anti-RBD IgG and IgM quantified by ELISA or (B) with the level of anti-SARS-CoV-2 S antibodies quantified by flow cytometry. Statistical significance was tested using Spearman rank correlation tests. (B) Indirect ELISA was performed using recombinant SARS-CoV-2 RBD and incubated with samples from COVID-19 negative or COVID-19 positive patients at different times after symptoms onset (T1, T2, T3, T4, Convalescent). Anti-RBD binding was detected using (B) anti-IgA-HRP. Relative light units (RLU) obtained with BSA (negative control) were subtracted and further normalized to the signal obtained with the anti-RBD CR3022 mAb present in each plate. Undetectable measures are represented as white symbols and limits of detection are plotted. Statistical significance was tested using Kruskal-Wallis tests with a Dunn’s post-test (* P < 0.05; ** P < 0.01; **** P < 0.0001). (C) The levels of anti-RBD IgA were correlated with the neutralization ID_50_ with SARS-CoV-2 S, the level of anti-SARS-CoV-2 S antibodies quantified by flow cytometry and the levels of anti-RBD IgG and IgM quantified by ELISA. Statistical significance was tested using Spearman rank correlation tests.

**Supplemental Figure 5.**
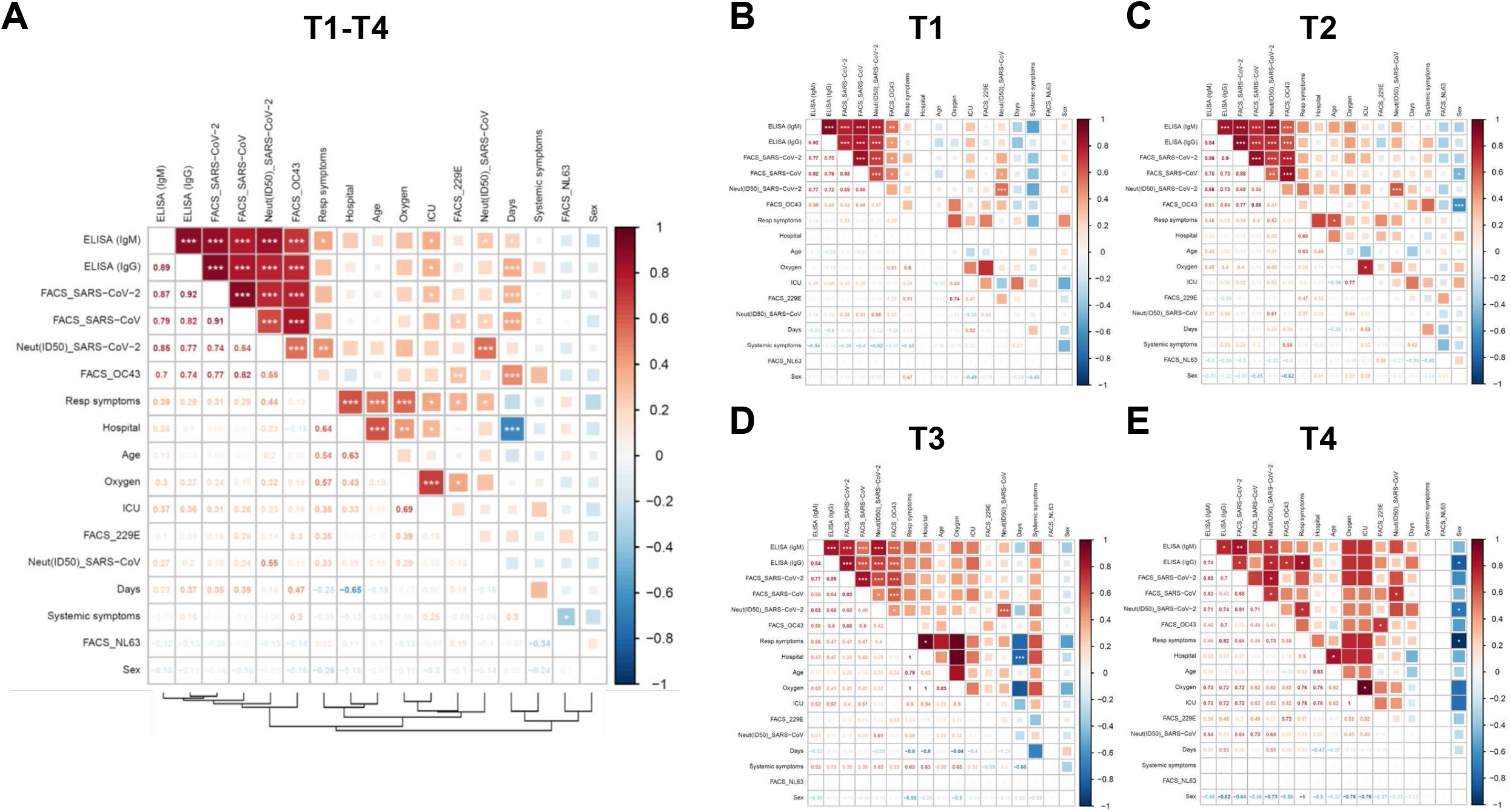
Correlations between serological measurements and clinical outcome. Correlograms were generated by plotting together all serological and clinical data obtained from acutely infected COVID-19+ patients (T1, T2 and T3), separated by time points (a-c) or all together (d) or using data obtained from convalescent patients (e). Squares are color-coded according to the magnitude of the correlation coefficient (r) and the square dimensions are inversely proportional with the P-values. Red squares represent a positive correlation between two variables and blue squares present negative correlations. Asterisks indicate all statistically significant correlations (*P < 0.05, **P < 0.01, ***P < 0.005). (a-e) Correlation analysis was done using nonparametric Spearman rank tests.

**Supplemental Figure 6.**
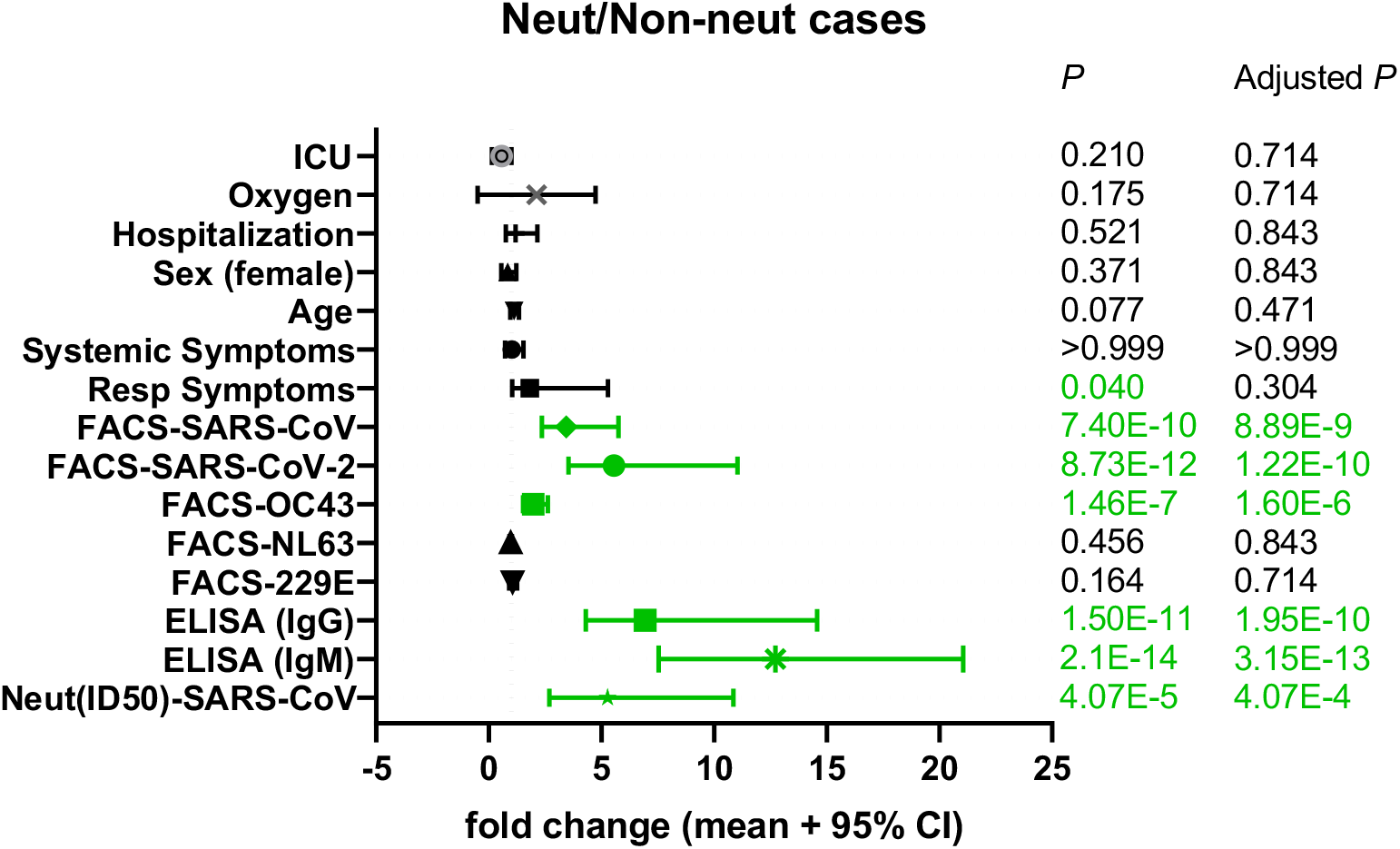
Clinical, demographic, and humoral factors associated with increased SARS-CoV-2 neutralization. Forrest plot of the association of SARS-CoV-2 neutralization with selected clinical, demographic, and humoral parameters. The fold change (mean and 95% confidence interval) of the parameters, listed on the y-axis, between neutralizers (ID_50_ >100) and non-neutralizers (ID_50_ <100) is displayed on the x-axis. Significance P and adjusted P values (Holm-Sidak method) are shown in columns to the right. Results with P<0.05 are highlighted in green.

## Notes

### Competing Interest Statement

The authors have declared no competing interest.

### Summary of Updates

Clinical information on the cohort was added and correlations with humoral responses were established. Additional cross-reactive assays were performed.

